# Detection of *Borrelia garinii* in the USA

**DOI:** 10.1101/2022.05.15.491491

**Authors:** Natalie Rudenko, Maryna Golovchenko, Ales Horak, Libor Grubhoffer, Emmanuel F. Mongodin, Claire M. Fraser, Weigang Qiu, Benjamin J. Luft, Richard G. Morgan, Sherwood R. Casjens, Steven E. Schutzer

**Author notes:** corresponding author: Biology Centre Czech Academy of Sciences, Institute of Parasitology, Branisovska 31, 37005, Ceske Budejovice, Czech Republic. Division of Lung Diseases, National Heart, Lung, and Blood Institute (NHLBI), National Institutes of Health (NIH), Bethesda, Maryland, USA.

## Abstract

*Borrelia garinii*, is a cause of Lyme disease in Europe and Asia. For the first time, we report it in the southeastern United States in rodents. Whole genome sequencing and phylogenetic analysis revealed that USA-located *B. garinii* is part of a clade consisting primarily of few European and majority of Far Eastern strains. Continued surveillance of wildlife hosts and ticks is necessary to assess the ecological status and public health risks of *B. garinii* in southeastern US.

## Introduction

Lyme disease (LD) is a multisystem disorder caused by *Borrelia burgdorferi* sensu lato (Bbsl, also known as *Borreliella*). *B. burgdorferi* sensu stricto, *Borrelia garinii* and *Borrelia afzelii* are responsible for most cases of LD worldwide (1, 2). *B. burgdorferi* is the only one of these three species that is normally found widely in North America, although *B. garinii* has been identified on islands off the coast of Newfoundland and Labrador, Canada (3-5).

We describe here the isolation and characterization of *B. garinii* from rodent hosts in South Carolina (USA) and provide the first report of the detection of *B. garinii* in the United States.

## Methods

### *Borrelia:* sources, cultivation and analyses

The two *Borrelia* isolates described herein, SCCH-7 and SCGT-19, were isolated from ear biopsies from a cotton mouse (*Peromyscus gossypinus*) and an eastern woodrat (*Neotoma floridana*) trapped in Charleston County, South Carolina in 1995 and in Georgetown County, South Carolina in 1996, respectively. *Borrelia* culture in Barbour-Stoenner-Kelly H medium, DNA purification and PCR analyses were as described (6). Initial PCR that detected the presence of coinfection was performed on the 5S-23S intergenic region (7) and was confirmed by species-specific PCR with primers designed on the basis of *ospA* (8). MLST analysis of eight housekeeping genes (*clpA, clpX, nifS, pepX, pyrG, recG, rplB* and *uvrA*) was performed as described (9).

### Isolation of *B. garinii* clonal cultures

Cultures in which the presence of *B. garinii* was confirmed were plated on solid medium according to a modified protocol (10) (Appendix Methods).

### Whole genome sequencing and genome assembly

Genome sequencing was performed using the Pacific Biosciences Sequel II system. Genome assembly was performed with the “Genome Assembly” tool in PacBio SMRTLink 10.2 using 150 Mb of the HiFi reads greater than 5kb (Appendix Methods).

### Nucleotide sequence accession numbers

Sequences have been deposited into GenBank. The genome assembly of SCCH-7 has been deposited to GenBank under BioProject PRJNA431102 with the BioSample accession SAMN26226110 (Appendix Results).

### Phylogenetic analysis

The maximum likelihood phylogeny of *B. garinii* strain from the concatenated dataset of eight housekeeping loci (184 isolates in total; 4791 nucleotides) was inferred in RAxML (Appendix ref. 3) under the GTR+G4 model. Phylogeographic analysis of diffusion on discrete space as implemented in BEAST (Appendix refs. 4, 5) was performed under the constant-size coalescent tree prior, and symmetric substitution model with BSSVS enforced (Appendix Methods).

### Comparison of closely related *B. garinii* genomes

Upon the discovery that the two USA *B. garinii* isolates (SCCH-7 and SCGT-19) were closely related to *B. garinii* type strain 20047, we proceeded to compare the chromosomal sequences among these isolates. We used BLASTN (11) to extract the sequences at the eight housekeeping loci (above) from the chromosomal sequence of SCCH-7 and the two released chromosomal sequences of the strain 20047 (CP018744 and CP028861, both unpublished). These gene sequences were subsequently concatenated for each genome and integrated into the MLST alignment. A phylogenetic tree was derived using IQTREE (12) with default parameters from an MLST alignment of 34 *B. garinii* isolates most closely related to the two USA isolates. To compare sequences for the entire chromosome, we aligned the SCCH-7 chromosomal sequence with the two published chromosomal sequences of the strain 20047 respectively using NUCMER (13).

## Results

### *B. garinii* from South Carolina rodents

The two *B. garinii* isolates discussed herein were cultured from the ear biopsy samples of two rodents from the southeastern United States: a cotton mouse, *Peromyscus gossypinus* (isolate SCCH-7) and an eastern woodrat *Neotoma floridana*, (isolate SCGT-19), both trapped in South Carolina. These cultures were part of a southeastern *Borrelia* culture collection of approximately 300 *Borrelia* isolates in the James H. Oliver, Jr. Institute of Arthropodology and Parasitology, Georgia Southern University, between 1991 and 1999. The presence of multiple Bbsl species in cultures was confirmed by PCR amplification from total DNA with a 5S-23S rRNA set of primers (7). Cloning of total PCR products into the pCR4-TOPO TA vector and sequencing of individual recombinants revealed multiple *Borrelia* species in numerous cultures, often present as co-infection, including *B. burgdorferi* sensu stricto (14), *B. bissettiae* (Rudenko, Golovchenko, Oliver Jr., unpublished), *B. kurtenbachii* (Rudenko, Golovchenko, Oliver Jr., unpublished), *B. carolinensis* (15) and *B. americana* (16). Unexpectedly, sequences with high similarity to *B. garinii* were detected in five cultures which were then plated on solid medium to separate the present species. Monoclonal populations of *B. garinii* were obtained from two of the five cultures. Two *B. garinii* positive clones, SCCH-7 clone 138 and SCGT-19 clone 19, were selected for further analysis.

### Whole genome sequence of *B. garinii* isolate SCCH-7

Isolate SCCH-7 was chosen for more detailed analysis and its whole genome sequence (WGS) was determined by single-molecule real-time (SMRT) PacBio methods (Appendix Methods). Like other Bbsl genomes (17-19), the SCCH-7 genome contains a linear chromosome and several linear (lp) and circular (cp) plasmids. It carries lp17, lp28-7, lp32-10, lp36 and lp54 linear plasmids and cp26, cp32-3 and cp32-6 circular plasmids (see 20 for discussion of plasmid types). The plasmid SCCH-7 sequences are typical of known *B. garinii* genomes and are also similar to those of strain 20047, although 20047 carries an lp28-4 plasmid that is lacking in SCCH-7. The SCCH-7 genome is 1,161,212 bp long (chromosome 906,106 bp, linear plasmids 168,083 bp, circular plasmids 87,023 bp). The linear chromosome and plasmid sequences include all telomeres, so this genome joins *B. burgdorferi* B31 and *B. mayonii* MN14-1539 genomes in being truly complete (21-23). The SCCH-7 chromosome differs from that of *B. garinii* strain 20047 by two single nucleotide variations (SNVs) and two indels from the sequence in CP028861 and by 8 SNVs and 4 indels from the sequence in CP018744; these are two 20047 chromosomal sequences produced by two independent research groups, S. Bontemps-Gallo and G. Margos et al., Bioproject PRJNA224116. Its plasmids were described briefly in Casjens et al. (19). This whole genome sequence demonstrates that SCCH-7 is a *B. garinii* isolate.

### Phylogenetic analysis

To understand the relationship of SCCH-7 and SCGT-19 to other *B. garinii* isolates, we PCR amplified and determined SCGT-19 sequences for the eight genes previously used in MLST analyses of Bbsl isolates, and we extracted these sequences from the SCCH-7 and 20047 whole genome sequences (WGSs) (4791 nucleotides *–* genes *clpA, clpX, nifS, pepX, pyrG, recG, rplB* and *uvrA* (9)). Figure S1 shows a phylogenetic tree of all isolates in this portion of the Bbsl species group, and Figure 1 shows a maximum likelihood (RAxML (Appendix, ref. 3)) tree of these MLST sequences that includes isolates SCCH-7 and SCGT-19 and the 178 other non-recurring isolates currently available of the closely related species *B. garinii* and *B. bavariensis*, as well as five isolates of *Borrelia turdi* as an outgroup (Table S1). Apart from one unusual isolate from European Russia (pubMLST ID:2488 “Om16-103-Iapr”) that is a sister branch to all the other isolates, the remaining 178 isolates form two clades of 39 and 139 isolates that agree with the previously defined *B. bavariensis* and *B. garinii* species, respectively (Figure 1) (24). The *B. bavariensis* group contains four European isolates and one Canadian isolate interspersed among a majority of Asian isolates suggesting that this is an Asian clade with several independent introgressions of western hemisphere types (25). The *B. garinii* clade is split into two major clades in agreement with Takano et al. (24). The larger one (108 isolates) includes 76 European isolates, 13 from continental Asia, 2 from Japan, 9 from Canada (Newfoundland and Labrador) and 8 from Iceland that are mostly distributed within this branch without an apparent pattern. The smaller *B. garinii* clade (31 isolates) indicated as “Asian” in Figure 1 contains 21 isolates from continental Asia and Japan, eight from Europe, and the two from the United States described here. The two USA isolates form a nested subclade with five strains of European origin in the “Asian” clade (Figure 1B). We also note that the Canadian *B. garinii’s* are in the “European” clade in Figure 1 and are not closely related to the USA isolates.

**Figure 1.**
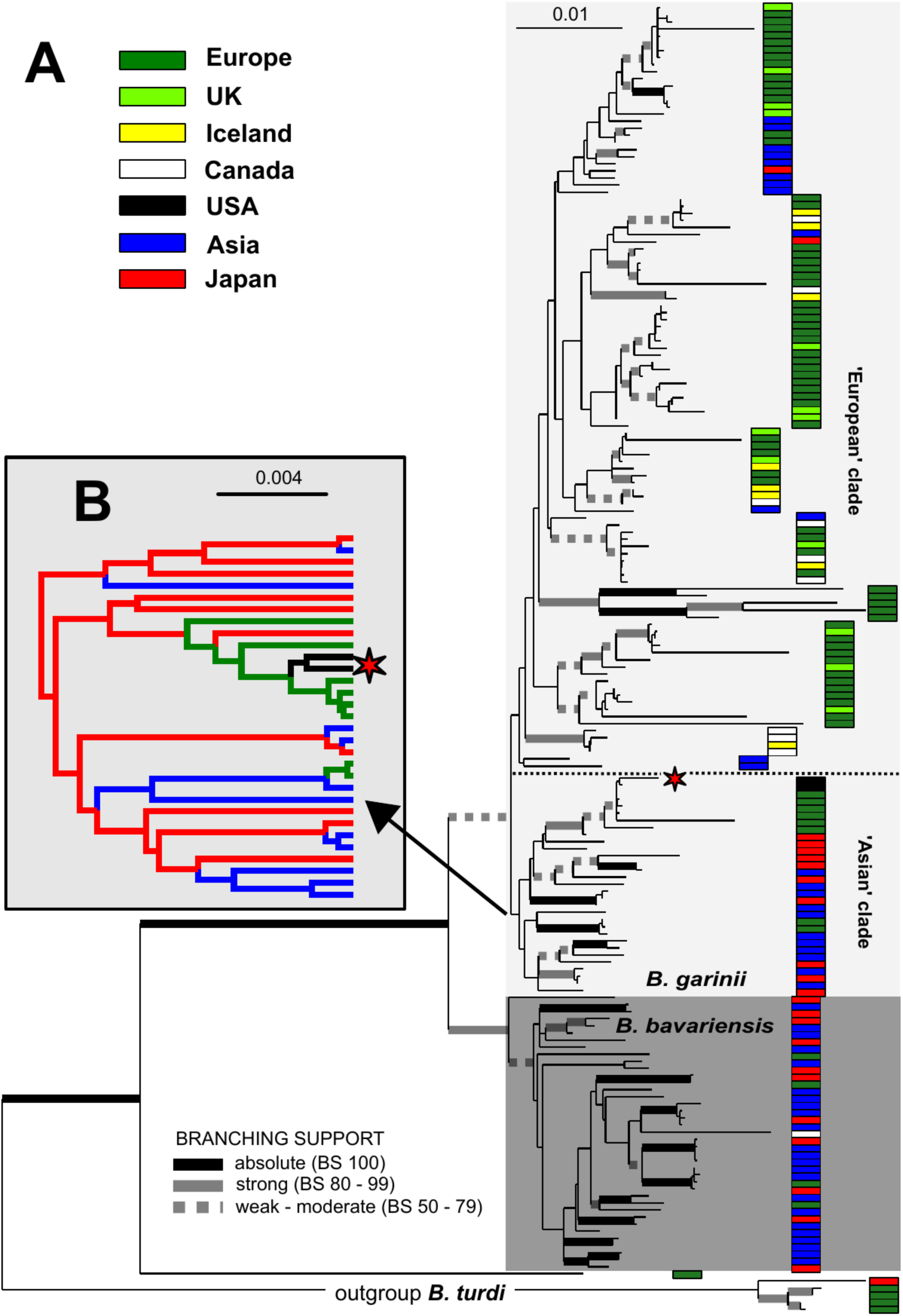
Phylogenetic trees of *B. garinii* and *B. bavariensis* strains. **A:** Maximum likelihood phylogeny of *B. garinii/B. bavariensis* rooted with *B. turdi*. The topology is based on the analysis of the partitioned dataset of eight ‘MLST’ genotyping loci (see Appendix Methods for details) under the GTR+G4 model (for each partition) in RAxML 8. The final alignment comprises of 184 taxa and 4791 nucleotide positions. The thickened branches represent the branching support as estimated by the non-parametric bootstrap analysis based on 1000 replicates in RAxML 8. For the sake of better readability, the support is categorized according to the scheme shown at the bottom of the tree. The isolates were clustered into seven categories according to their geographic origin, which is color-coded according the scheme in the upper-right part of the tree on the topology. The position of two US isolates is indicated by an asterisk. **B:** Subset of results phylogeographic analysis of diffusion on the discrete space showing the estimated geographic origin of the inner branches for the ‘Asian’ clade of *B. garinii*. The full topology is shown in Figure S2. For the full details on the method see the relevant part of Appendix Methods.

To shed more light on the origin of the two USA isolates and the evolutionary history of *B. garinii* in general, we performed phylogeographic analysis of diffusion in discrete space as implemented in BEAST (Appendix refs. 4, 5). This Bayesian method enables each node of the tree to predict the ancestral state of a given discrete trait (in this case the geographic origin of the strain) and to map it on the resulting topology. The resulting tree topology (Figure S2) as inferred under “Coalescent: constant size,” separated the isolates into a number of groups. “*Bavariensis*” and “Asian” clades, whose composition corresponds to the MLST topology in Figure 1, are both predicted to originate in Japan. As they do in Figure 1, the two USA isolates reside in the Figure S2 “Asian” clade, nested among few sequences from Europe. The “European” isolates are split into three clades, the first of which is found at the base of the whole *B. bavariensis/B. garinii* portion of the tree. It is composed of five divergent *B. garinii* isolates from Slovakia and may be a result of long-branch-attraction phylogenetic artifact caused by employing the suboptimal phylogenetic model available for the BEAST software (26). This clade is separated from the remaining “European” clades by the “Asian” clade, whose position is shifted compared to the Figure 1 tree. The terminal position of the European strains in this clade suggests a secondary introduction into Europe.

The affiliation of USA isolates with Europe and the broader “Asian” *B. garinii* clade is consistently shown in both trees (Figures 1 and S2) and is supported by phylogeographic reconstruction using the BSSVS algorithm (Appendix refs. 4, 5). However, due to the large number of isolates and relatively low number of phylogenetic-informative positions (*i*.*e*., high sequence similarity) in our dataset, the bootstrap support of inner branches was not high. Therefore, we tested the independent evolutionary history of USA and Canadian isolates using the approximately-unbiased (au) topology test (Appendix, ref. 7). First, we force-constrained the monophyly of two USA isolates with each of the nine Canadian isolates; we then let RAxML re-optimize the general topology. The per-site log likelihood scores of those alternative topologies were then compared to the original MLST topology (Figure 1) using the au-test in Consel (Appendix, ref. 8). The resulting *p*-values (ranging from 1.48e-36 to 1.9 e-2, see also Table S2) support the rejection of a common origin of Canadian and USA *B. garinii*. We conclude that, in contrast to the Canadian isolates which were found on islands in the Atlantic and may have been introduced there from Europe and/or Iceland by seabirds (27), *B. garinii* may have arrived in the USA from the Far East (Japan) by way of Europe.

### Chromosomal relationships with closely related *B. garinii* genomes

The maximum likelihood tree of 32 closely related *B. garinii* genomes based on the eight housekeeping loci (Figure S3) showed that the two USA isolates are a part of a “European” clade, which in turn is a part of a mostly Asian group (Figure 1). Table S3 lists all sequence differences at the eight housekeeping loci among the strains, most closely related to the two USA isolates. Consistent with the phylogenetic tree (Figure 1), the genome-derived MLST SCCH-7 sequence and two independent 20047 sequences are almost identical; curiously the reported 20047 MSLT sequence has two differences in the *recG* gene compared to the above three WGSs (this is likely due to sequencing errors). The above sequence identities strongly support a recent common European origin of the USA isolate SCCH-7.

In contrast to SCCH-7, strain SCGT-19 shows a distinct MLST haplotype defined by 16 single nucleotide variations (SNVs) and one short indel (Table S3). The SCGT-19 versions of some of these SNVs and the indel are found in other *B. garinii* strains from Japan and Europe, suggesting that they are unlikely to be sequencing errors. Further, consecutive runs of SNVs at the *clpA* and *clpX* loci strongly indicate that their origins are due to recombination and not to *de novo* mutation. The differences between SCCH-7 and SCGT-19 suggests that a migration or importation of *B. garinii* from Eurasia to the USA may have consisted of multiple strains of a source population.

## Discussion

Our results provide strong genomic evidence that *B. garinii* has appeared in rodents in the state of South Carolina. This is the first report of *B. garinii* in the USA. Specifically, 5 rodent samples tested positive for *B. garinii*, and from these two independent *B. garinii* cultures, SCCH-7 clone 138 and SCCH-19 clone 19, were propagated. MLST analyses of both USA isolates and whole genome sequencing of SCCH-7 showed that they are very closely related to a subset of European and Asian isolates and not closely related to the Canadian *B. garinii* strains. How it came there or when remains unknown despite intriguing comparative genomic data. The potential clinical impact requires observation and case surveillance in humans. Serologic studies in humans and the rodents from the area would be helpful information to assess clinical risk and could complement tick analysis from the area. Even though nearly 25 years has passed after the isolates were obtained, no cases in humans in the region have been reported.

It is of value to characterize the mechanisms of spread of infectious diseases for effective control measures. This should include migration of infected vertebrate (28, 29). Given the report of *B. garinii* in seabird colonies on islands in far eastern Canada (3-5, 30), one might think it could spread south to the US, but it has not. This is in the context that birds can function both as biological carriers of Borrelia spirochetes and as transporters of attached ticks infected with *Borrelia* (30-32). *Ixodes scapularis* has been confirmed to be able to transmit *B. garinii* (33, 34).

A plausible, but still speculative, cause of *B. garinii* detection in South Carolina could be related to ships that travel worldwide and infested with mice and rats (35). This would be stronger if the rodents were of the same types or at least known to transmit *B. garinii* from one species to the ones in South Carolina.

The public health importance of our findings will depend on the pathogenic potential of the *B. garinii* strains detected in South Carolina. Since nearly a decade has past multi-year longitudinal studies are needed to determine if the two isolates reported here represent a small and transient population, or an emergent and growing *B. garinii* population in the southeast US. Analysis, if the US *B. garinii* strains are pathogenic to humans, should be the topic of further investigation as new *B. garinii* strains are introduced in environments where humans are at risk of acquiring the infection (36-38).

## Appendix

- Figure S1. Maximum likelihood phylogeny of *Borrelia burgdorferi sensu lato* (*Bbsl*) based on eight MLST loci
- Figure S2. Maximum likelihood phylogeny of 32 *B. garinii* closely related to the two US isolates (SCCH-7 and SCGT-19)
- Table S1. *Bbsl* strains included in the phylogenetic study
- Table S2. Result of approximately-unbiased topology test
- Table S3. Single-nucleotide variants among strains closely related to the two US *B. garinii* isolates.

## Acknowledgements

Supported in part by AI139782 (WQ) from the National Institute of Allergy and Infectious Diseases (NIAID) of the US National Institutes of Health (NIH); Biology Centre, Institute of Parasitology Czech Academy of Sciences (NR, MG, AH, LG); by Institute for Genomic Sciences (CF, EM), by New England BioLabs (RM); Steven and Alexandra Cohen Foundation and the Global Lyme Alliance (BL).

We acknowledge all former members of the James H. Oliver, Jr. Institute of Arthropodology and Parasitology, Statesboro, Georgia, USA, who contributed to the establishment and development of the southeastern spirochete collection during 1991–1999.

## Disclaimers

The authors declare no conflict of interest. The opinions expressed by authors contributing to this journal do not necessarily reflect the opinions of the institutions with which the authors are affiliated. Emmanuel Mongodin, PhD, contributed to this work as an employee of the University of Maryland School of Medicine. The views expressed in this manuscript are his own and do not necessarily represent the views of the National Institutes of Health or the United States Government.

## About The Author

Dr. Rudenko holds a PhD degree in molecular and cellular biology and genetics and is a senior researcher at the Institute of Parasitology, Biology Centre Czech Academy of Sciences, Czech republic. Her primary research interests include spirochetes from *Borrelia burgdorferi* sl complex and ecological, epidemiological, and molecular aspects of Lyme disease.

## Detection of *Borrelia garinii* in the USA

### Appendix

#### Supplementary Methods and Results

##### Isolation of *B. garinii* clonal cultures

Colonies that appeared 5 -7 days after plating were re-cultured in liquid BSK-H medium, and PCR analysis of monoclonal cultures was performed as described above. All reactions were prepared in a dedicated area, and precautions were taken to avoid contamination of supplies, equipment and employees’ personal safety items in pre-and postamplification activities. Negative (no template) and positive controls (*B. burgdorferi* B31 DNA) were present in each amplification series. DNA of *B. garinii* was not used in any step of the PCR amplifications, and there were no *B. garinii* cultures in the Oliver laboratory. Sanger type sequencing of each amplicon was conducted in both directions, with the same primers used for PCR.

##### Nucleotide sequence accession numbers

Sequences determined in this study have been deposited into GenBank with the following accession numbers for SCCH-7/SCGT-19: 5S-23S IGR–KP795350/KP795363; and 16S-23S ITR–KP795352/KT285872. The accession numbers for the eight housekeeping genes for SCCH-7/SCGT-19 are clpA, KP795360/KT285880; clpX, KP795358/KT285878; nifS, KP795357/KT285877; pepX, KP795359/KT285879; pyrG, KP795356/KT285876; recG, KP795355/KT285875; rplB, KP795353/KT285873; and uvrA, KP795354/KT285874. The genome assembly of SCCH-7 has been deposited to GenBank under BioProject PRJNA431102 with the BioSample accession SAMN26226110.

##### Whole genome sequencing and genome assembly

Genome sequencing was performed using the Pacific Biosciences Sequel II system. Total genomic was isolated from SCCH7 cells using DNeasy Blood & Tissue (Qiagen, Germany) according to the manufacturer’s instructions. The gDNA sample contained relatively short fragments mostly less than 5kb, so no shearing was performed. A random library was prepared using the PacBio SMRTbell express template kit 2.0 according to the manufacturer’s instructions. Sequencing was performed using one Sequel II cell which generated 178Gb total sequence and 1.37M High Fidelity (HiFi) reads totaling 2.4Gb with mean length of 1757bp and mean quality of QV60. To facilitate assembly the subset of HiFi reads longer than 5000bp was generated which yielded 210Mb sequence in 32.6k reads with mean quality of QV42.

Genome assembly was performed with the “Genome Assembly” tool in PacBio SMRTLink 10.2 using 150 Mb of the HiFi reads greater than 5kb (100X downsample with 1.5Mb expected genome size). The assembly was polished by the PacBio Arrow algorithm, and the telomere ends were examined manually by comparison to multiple individual HiFi reads.

##### Phylogenetic analysis

The sequences of eight housekeeping loci (*clpA, clpX, nifS, pepX, pyrG, recG, rplB, uvrA*) of *Borrelia* isolates were from the PubMLST database (https://pubmlst.org/), and sequences of several additional relevant isolates were added from our collection. The resulting dataset was aligned with the MUSCLE aligner (1) implemented in SeaView 4 (2). To ensure the frame integrity of codon information, we translated the sequences into amino acids first, then aligned and back-translated them into nucleotides. Phylogenetic analysis with the maximum likelihood method was performed in RAxML (3) under the General tie-reversible substitutional matrix, with nucleotide frequencies estimated from the dataset and four gamma-corrected rate site classes (GTR+G4+F).

The best topology, and the branching support values were calculated by using the rapid bootstrap inferences from 1000 replicates followed by a thorough ML search (‘-f a’ parameter) in RAxML. Phylogeographic analysis of diffusion on discrete space as implemented in BEAST (4,5) was performed under the GTR+G4 matrix, constant-size coalescent tree prior, and symmetric substitution model with BSSVS enforced. To make the analysis more feasible, we simplified the location coding into following broad geographic categories: United Kingdom, Continental Europe, Asia, Japan, USA, Canada, and Iceland. To obtain sufficient effective sample sizes for the estimated parameters, we ran Monte Carlo Markov Chains (MCMC) for 100 million generations subsampling every 10000 trees. The MCMC convergence was then inspected, and “burnin” value was selected in Tracer (6). The final topology, including the reconstruction of distribution of *B. garinii* strains was summarized in TreeAnnotator (4), and the topology was visualized in FigTree. The approximately unbiased test (7) as implemented in Consel (8) was used to test alternative phylogenetic topologies.

---

**Figure S1.**
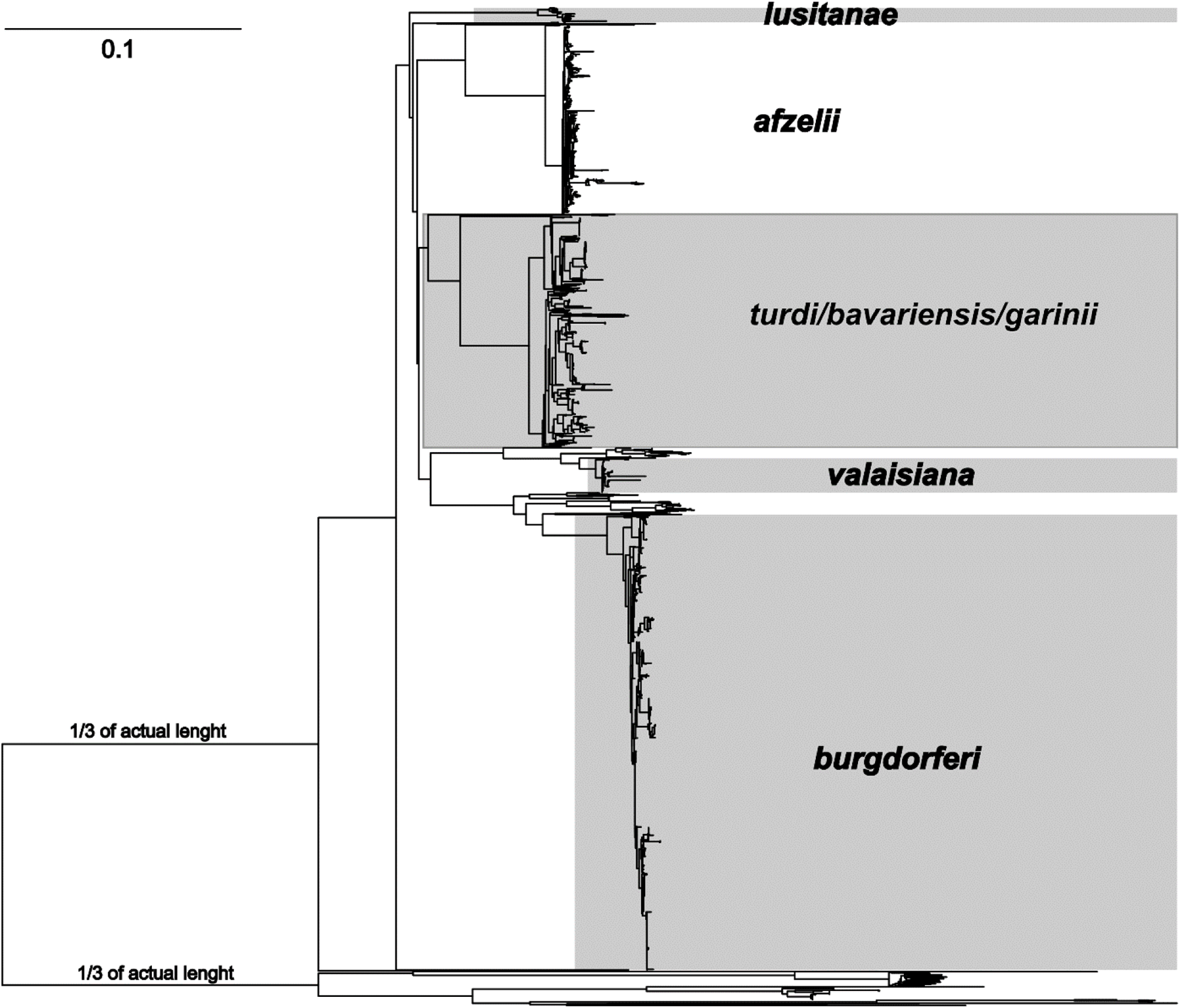
Unrooted maximum likelihood phylogeny of Borrelia based on the analysis of eight ‘MLST’ genotyping loci (see Appendix Methods for details) under the GTR+G4 model implemented in RAxML 8. The final alignment comprises of 2977 taxa and 4815 nucleotide positions. Please note that the length of basal branches was reduced from the formatting reasons.

**Figure S2.**
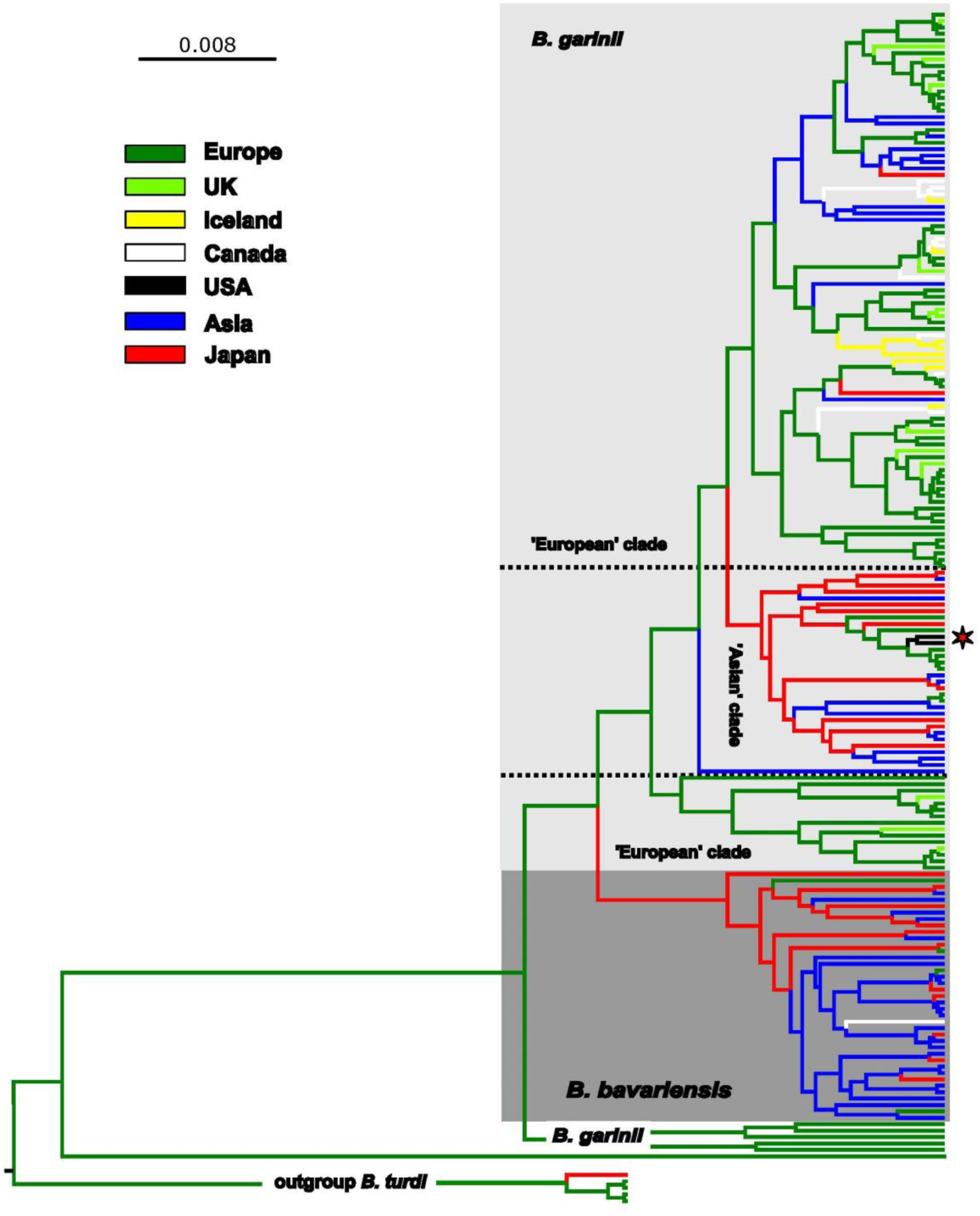
Phylogeographic analysis of *B. garinii/B. bavariensis* rooted with *B. turdi* using the diffusion on the discrete space algorithm implemented in BEAST under the GTR+G4 matrix, constant-size coalescent tree prior, and symmetric substitution model with BSSVS enforced. The isolates were clustered into seven categories according to their geographic origin, coded in form of branch colors according the scheme in the upper-right part of the tree on the topology. The position of two US isolates is indicated by an asterisk.

**Figure S3.**
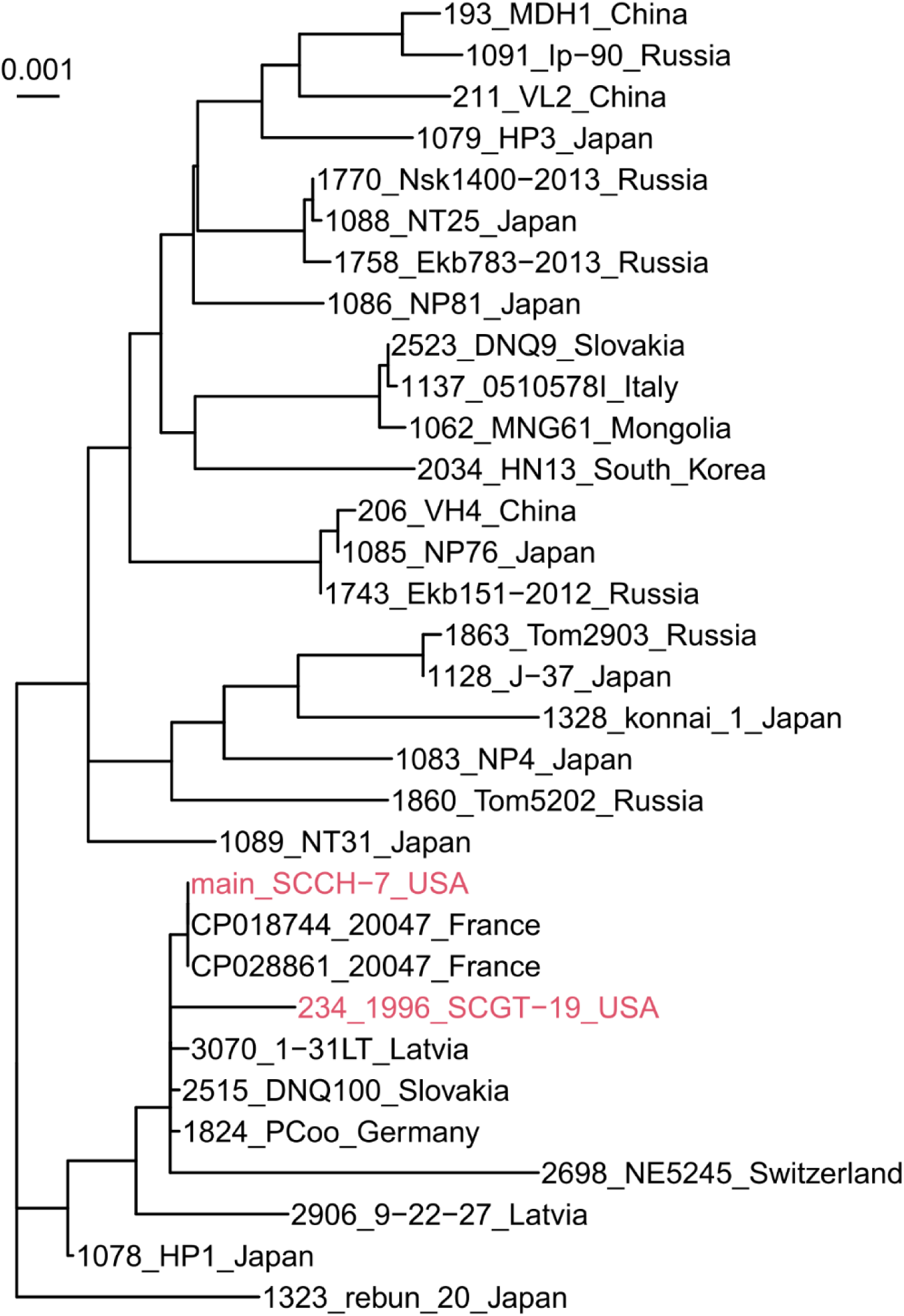
Maximum likelihood tree of 32 closely related *B. garinii* isolates based on sequences at the eight housekeeping loci. The maximum likelihood tree was inferred with IQTREE. All branches shown are supported by a bootstrap value of 80% or more. Two US isolates of *B. garinii* are highlighted in red. The SCCH-7 MLST sequence, grouped with the two previously sequenced genomes of the strain 20047 (CP028861 and CP018744) was derived from the genome sequence.

**Table S1.**
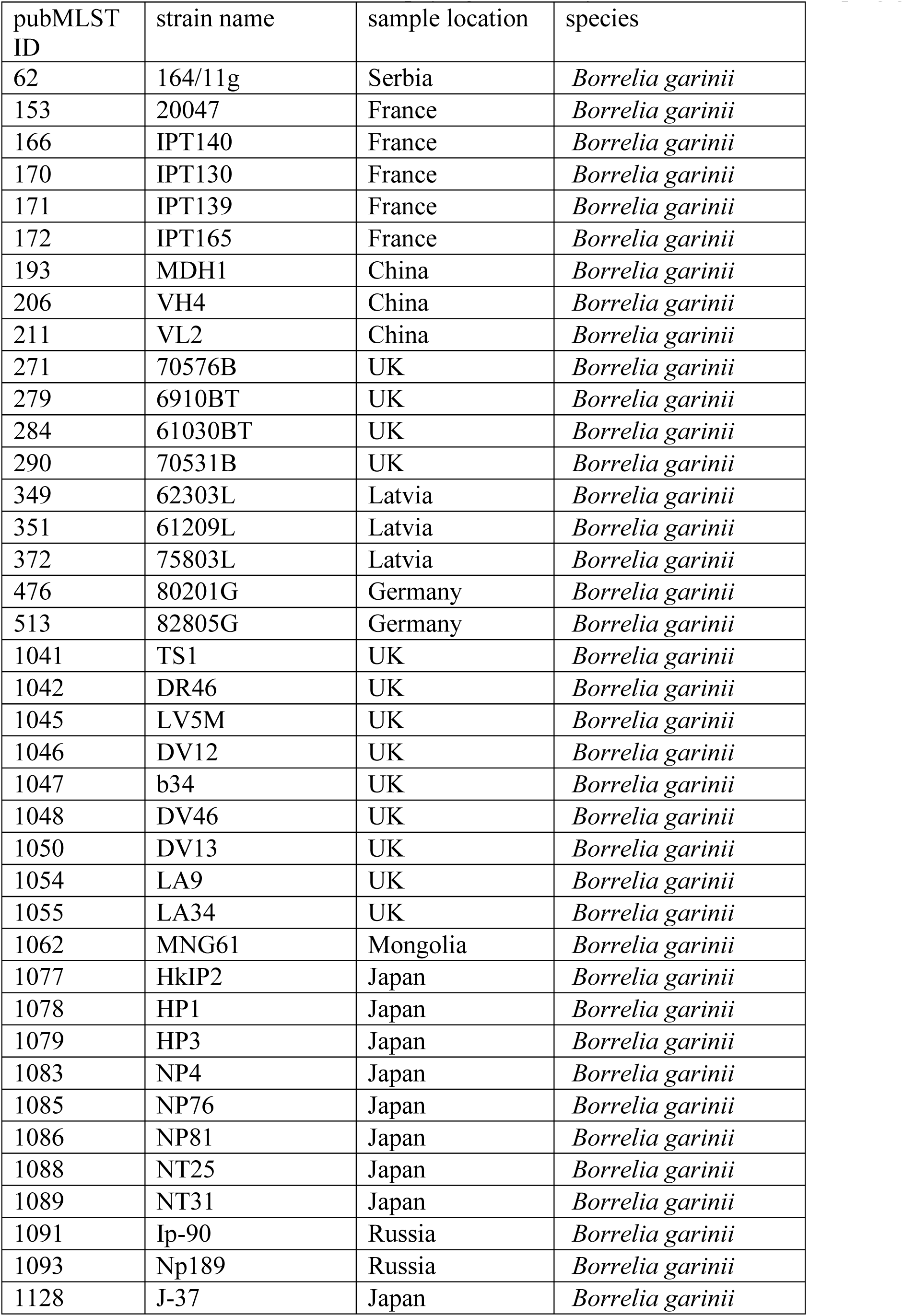
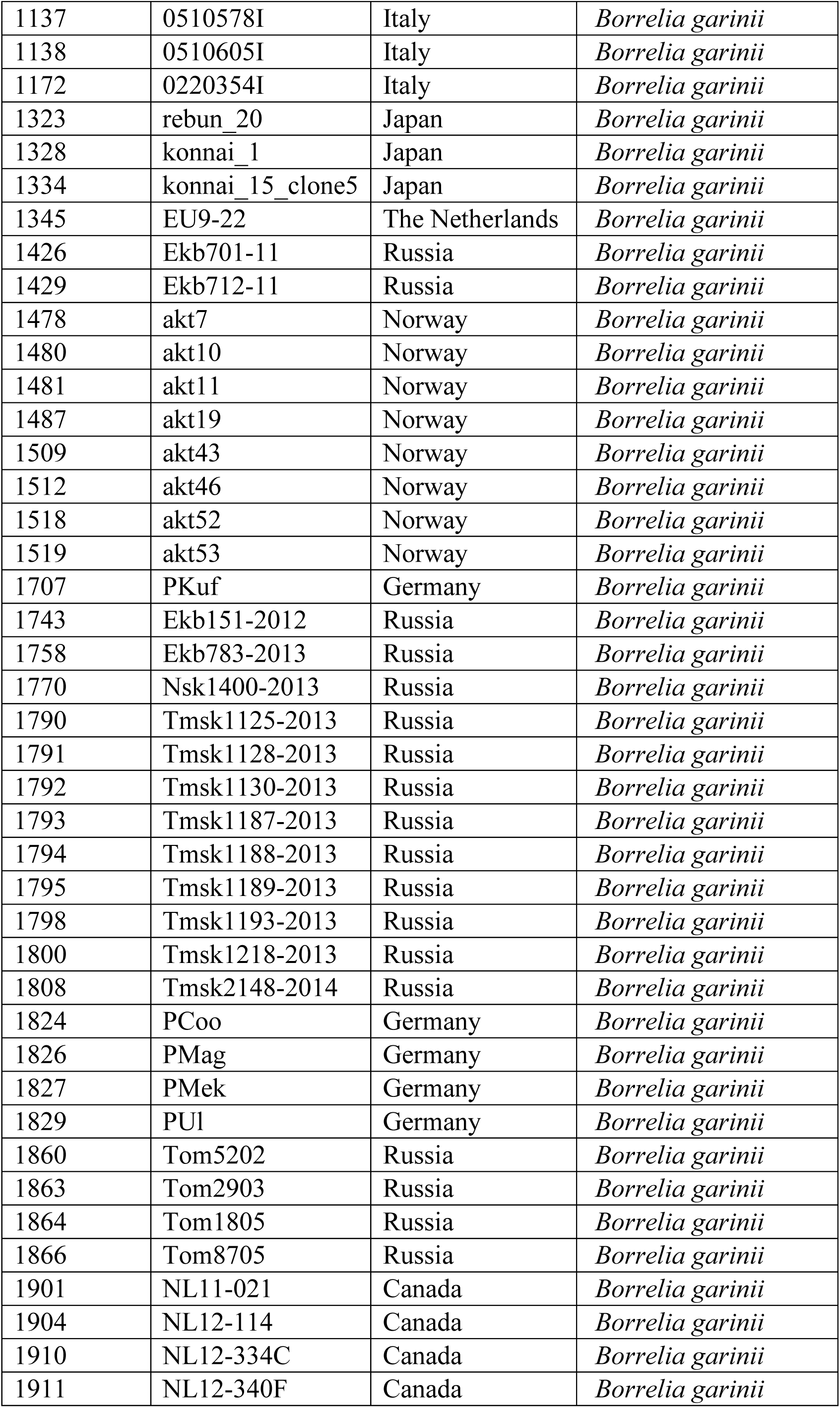
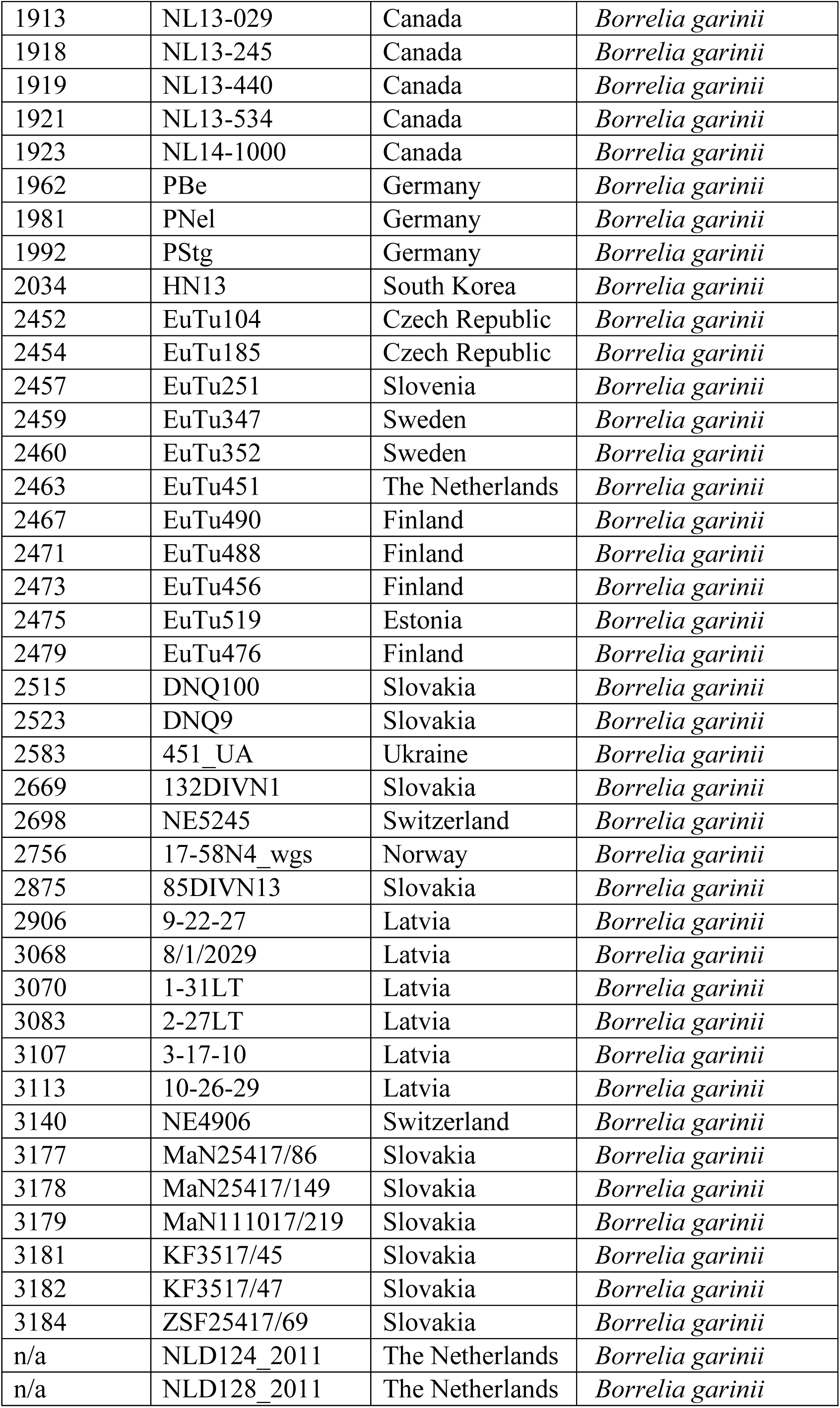
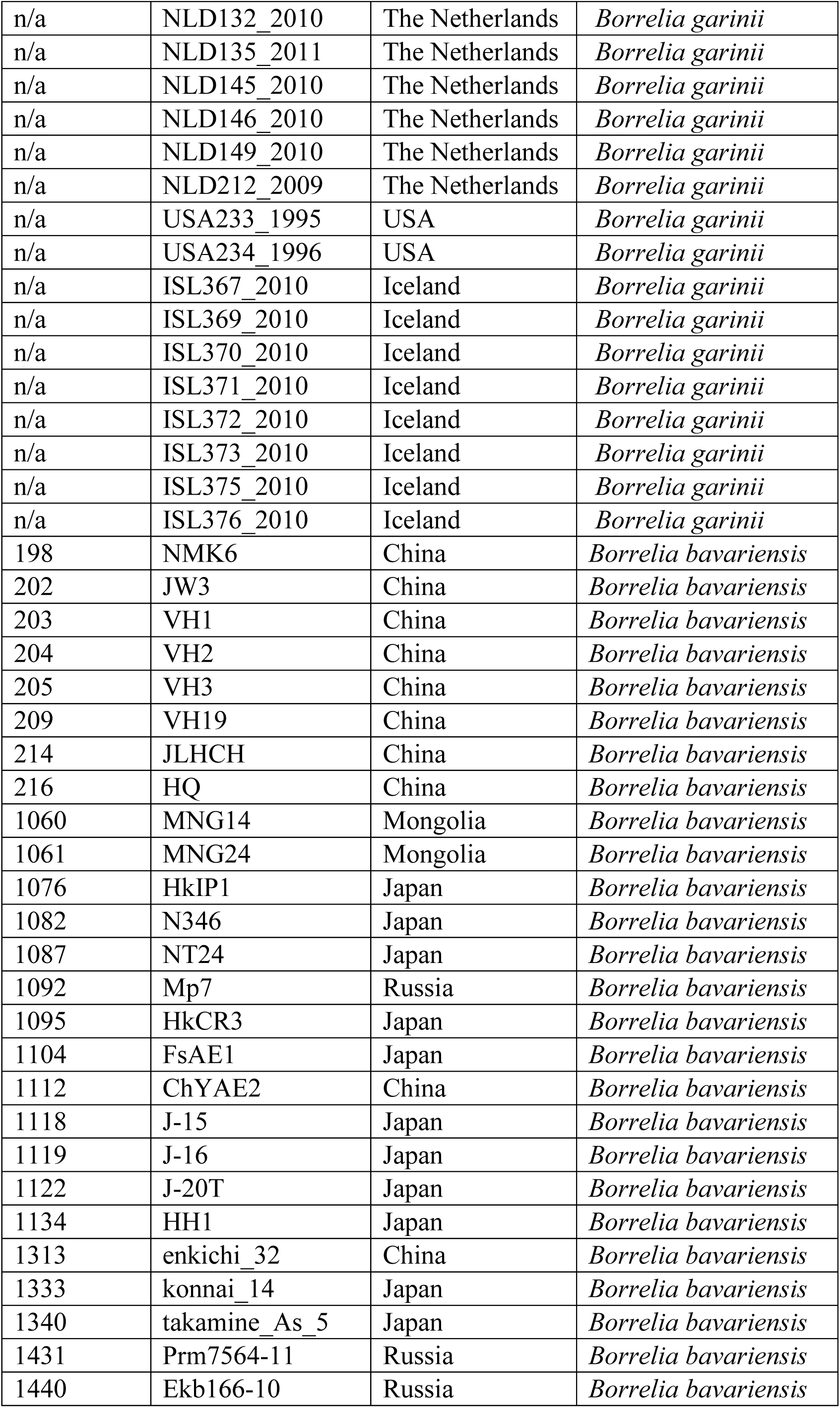
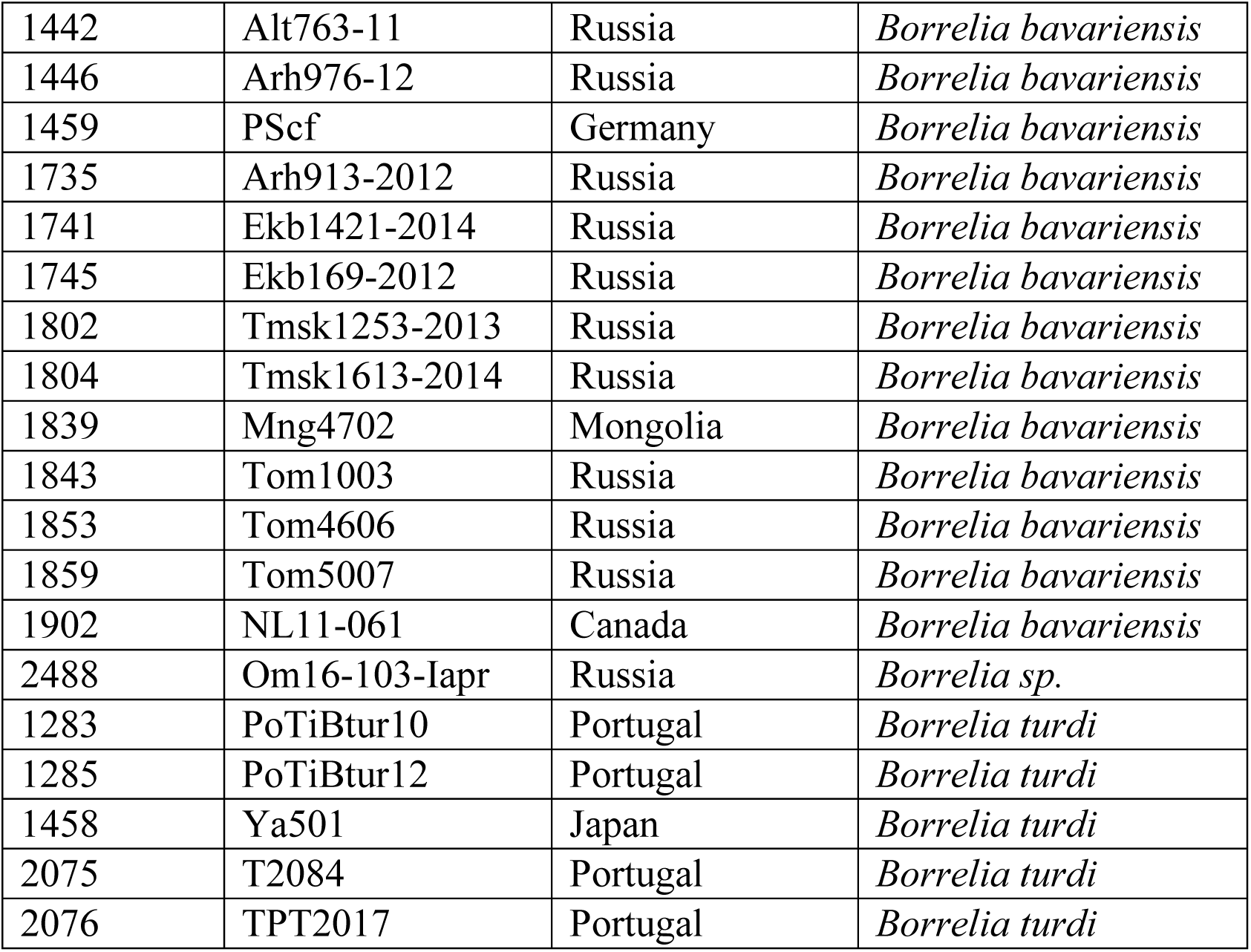
*Borrelia* strains included into phylogenetic analysis based on 8 housekeeping genes

**Table S2.** Results of Approximately-Unbiased (au) topology test comparing the original most-likely topology as seen on Fig 1 (row 1), with the alternative topologies with enforced Canadian-US B. garinii monophyly (rows 2-11). The p-AU column denotes the statistical signicance (p-values) of the test for individual topology.

| Tree | logL | deltaL | p-AU |
| --- | --- | --- | --- |
| 1 | -22917.7 | 0 | **0.995** |
| 2 | -23054.4 | 136.65 | 0.0002 |
| 3 | -23090.9 | 173.17 | 0.000748 |
| 4 | -23143.6 | 225.85 | 6.44E-05 |
| 5 | -23201.7 | 283.95 | 1.48E-36 |
| 6 | -23030.6 | 112.85 | 0.0192 |
| 7 | -23181.5 | 263.79 | 3.62E-07 |
| 8 | -23159.1 | 241.34 | 0.000139 |
| 9 | -23099.5 | 181.79 | 0.00125 |
| 10 | -23051.1 | 133.35 | 0.00791 |
| 11 | -23055.2 | 137.41 | 0.00345 |

**Table S3.**
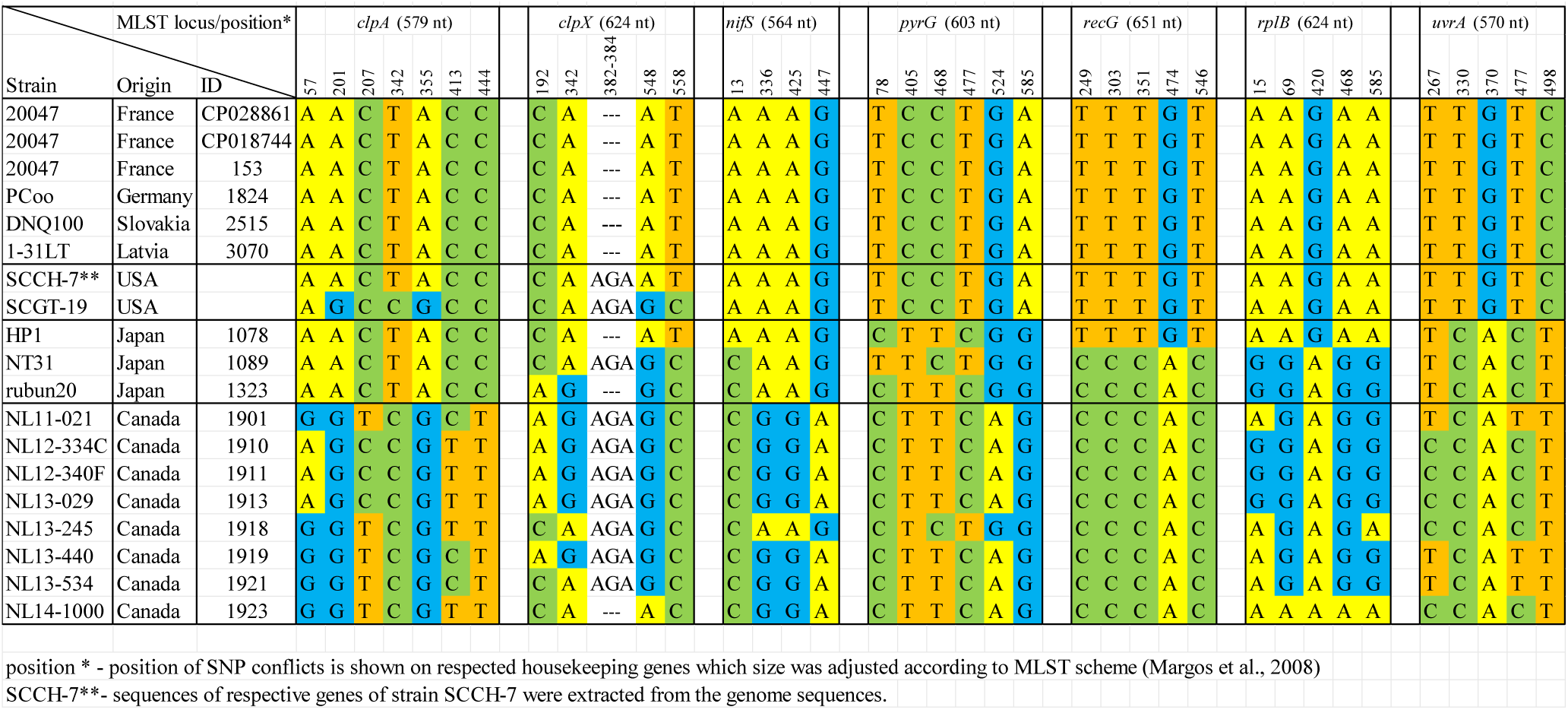
Distribution of single nucleotide variants (SNVs) among the housekeeping genes (MLST scheme).

